# Removable cranial window for sustained wide-field optical imaging in mouse neocortex

**DOI:** 10.1101/2020.01.14.905851

**Authors:** Susie S. Cha, Mark E. Bucklin, Xue Han

## Abstract

Attempts to image neocortical regions on the surface of mouse brain typically use a small glass disc attached to the cranial surface. This approach, however, is often challenged by progressive deterioration in optical quality and permits limited tissue access after its initial implantation. Here we describe a design and demonstrate a two-stage cranial implant device developed with a remarkably versatile material, polydimethylsiloxane, which facilitates longitudinal imaging experiments in mouse cortex. The system was designed considering biocompatibility and optical performance. This enabled us to achieve sustained periods of optical quality, extending beyond a year in some mice, and allows imaging at high spatio-temporal resolution using wide-field microscopy. Additionally, the two-part system, consisting of a fixed headplate with integrated neural access chamber and optical insert, allowed flexible access to the underlying tissue offering an expansive toolbox of neuromanipulation possibilities. Finally, we demonstrate the technical feasibility of rapid adaptation of the system to accommodate varying applications requiring long-term ability to visualize and access neural tissue. This capability will drastically reduce wasted time and resources for experiments of any duration, and will facilitate previously infeasible studies requiring long-term observation such as for research in aging or the progression chronic neurological disorders.

## Introduction

Technical advances in the tools available to visualize and manipulate dynamic cellular processes in real time have provided an unprecedented lens to probe the complex neural circuits in vivo (1–6). Using a wide-field fluorescence microscope with a scientific-CMOS camera, we can record activity in hundreds of distinct neurons across wide areas of the brain of awake behaving mice (7–9). Additionally, when combined with optical or pharmacological manipulation and/or electrophysiological recording, optical imaging provides a sophisticated approach to investigate neuronal function within the larger neural network (10–14).

One of the key advantages of in vivo imaging regards the ability to observe and record from the same brain region for extended periods to track long term changes (1, 9, 15, 16). This ability relies heavily on maintaining a clear optical light-path by forming a stable non-scattering optical interface with neural tissue overlying the targeted brain region. Attempts to image neocortical regions on the surface of mouse brain typically use a small glass disc attached to the cranial surface to seal and protect the craniotomy (16–19). This approach, however, is often challenged by progressive deterioration in optical quality (19–21). The degradation is observed as a cloudy layer that gradually overtakes the fluid filled gap between the cranial window and the brain tissue, and is thought to arise from the natural inflammatory response that follows a craniotomy (18–20, 22–24). As granulation tissue grows, its inhomogeneous structure scatters light at the brain-to-window interface, which consequently degrades optical quality and blurs fluorescence signals.

Efforts to overcome this problem by adding purely mechanical features to the cranial window have involved attaching spacers made of agarose (18, 25), silicone (20, 21, 26, 27) and glass (16) to the window’s brain-facing surface that compensate for the thickness of removed bone. These approaches report delaying tissue regrowth for up to a few months before optical quality deteriorates. These modest results indicate a valid basis underlying this approach and suggest that extending this strategy by starting with a design and material not limited by the fixed form of flat glass optical windows could yield some improvement. Additional elements of a chronic cranial imaging window intended to mitigate degradation by granulation tissue typically target inflammation, the primary source stimulating the process. These include the aseptic design of seals and features (28), selective use of biocompatible materials (10, 20, 21, 23, 29), and perioperative administration of anti-inflammatory and antibiotic drugs (16, 30, 31). While these designs have improved longevity, they remain limited in terms of long-term access to the cortical tissue, post-installation. The ability to access and manipulate tissue during real-time imaging opens the door in experimental designs to an expansive toolbox of neuromanipulation possibilities allowing exploration of uncharted connectivity and dynamic processes of the brain (17). Several strategies have been reported to gain access to regions below glass cranial windows by incorporating features such as an access port sealed with elastomer (12, 13), infusion cannula (14, 32), or the use of microfluidic channels (25). Nonetheless, the approaches limit the tissue accessibility to a single designated site predetermined before an experiment begins and do not offer uniform access over the imaging area.

To address the relative restrictions using glass as cranial windows, a number of alternative approaches have highlighted the use of silicone elastomer for cranial windows (10–12, 33, 34). For example, polydimethylsiloxane (PDMS) is optically clear, non-toxic and chemically inert and can be molded to take any shape or exhibit any desired feature, not necessarily sacrificing the imaging field of the window. These properties combine to offer a remarkably versatile material, particularly favorable for prototype development for projects with demanding specifications for biocompatibility and optical performance. A well-known and widely used example is the artificial dura for in vivo optical imaging in non-human primates (10, 11, 24, 35). This chronic implant device is placed in and covers a craniotomy and sits protected within a chronic cranial recording chamber. It mitigates tissue regrowth, and interfaces with a cylindrical insert – also made of PDMS – for optical imaging of neocortex. Additionally, the artificial dura is thin enough to enable access to underlying tissue for penetrating electrodes, which penetrate easily and leave a tight seal after withdrawal. Yet the efforts for translating this design windows for small research animals using silicone elastomer have thus far lacks the history of exploration it deserves (7, 12, 33, 34). And a system with long-lasting optical quality and flexible tissue accessibility remains to be developed or explored for rodent models.

In this paper, we describe a design and demonstrate a two-stage cranial implant device, developed to facilitate longitudinal imaging experiments in mouse neocortex. The primary capability requirements for this design are:

1. Long-term stability of an optically clear light-path to cortical surface
2. Intermittent physical access to imaged region at any point in study

The system design considered biocompatibility and optical performance to facilitate integration in place of the removed bone flap, enabling us to achieve sustained periods of optical quality which extended beyond a year in some animals. The two-part system, consisting of a fixed headplate with integrated neural access chamber and optical insert, allowed flexible access to the underlying tissue. The utility of our design is demonstrated through chronic optical imaging of calcium dynamics in the cortex using a wide-field microscope and acute interventions to the tissue upon removal and replacement of the cranial window from the headplate. Finally, we demonstrate the technical feasibility of rapid adaptation of the system that can accommodate a variety of applications, further extending our ability to visualize and access neural tissue.

## Results

Here we report the design for a head-fixation and cranial window device, and the procedures for surgical attachment. The sections below describe the features of each component, and also report the critical elements that contribute to the performance and capabilities of our cranial implant device. The following sections provide a detailed report of the system performance observed during evaluation. The final section reports the adaptability of the system with the demonstration of the latest design.

### Cranial Window System

Many design features, and procedures for installation were introduced and developed to mitigate tissue growth for the sustained optical quality of the window. Other features were included to enhance imaging performance in awake behaving animals, or to facilitate repeatable localization of image fields across sessions and animal subjects.

The cranial implant device is composed of two parts: a head-plate with an integrated chamber, and an optical insert (Figure 1). The headplate is bonded to the dorsal surface of the animal’s skull. The optical insert – sometimes referred to as a “cranial window” – seals the chamber and establishes an optical interface with the animal’s brain through craniotomy sites in the chamber floor.

**Fig. 1.**
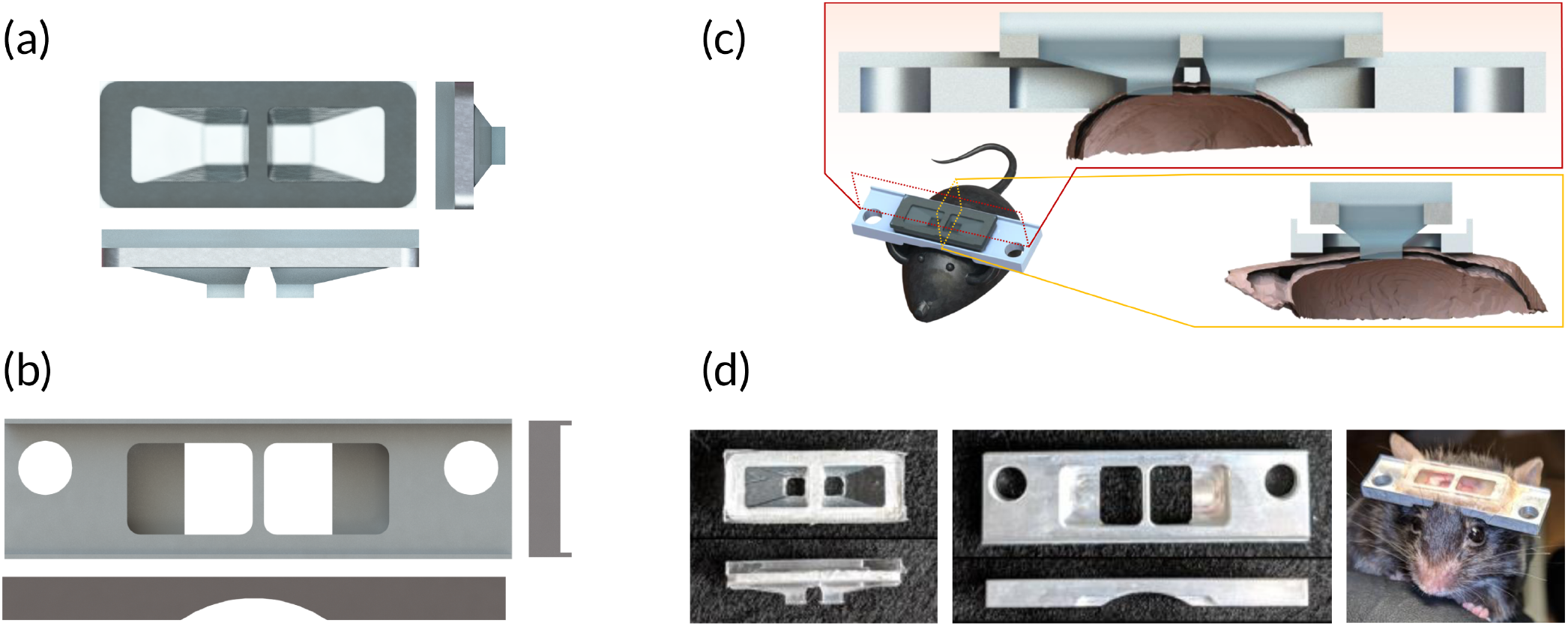
Multi-component cranial window system. (a) Rendered model of PDMS-based optical insert (also referred to as cranial window). The optical insert establishes an optical interface with the animals brain through craniotomy site. (b) Rendered model of headplate with an integrated chamber. The headplate is bonded to the mouse cranium and provides mechanically stable coupling between the animal skull and the headplate holder to reduce brain motion relative to the imaging system. (c) Cross-sectional view of the complete system as exists after surgical attachment to mouse cranium. The optical insert is attached to the headplate directly over the craniotomy and makes a gentle contact with the exposed brain tissue. (d) Matching photographs.

#### Headplate

The bottom surface of the headplate is curved to conform to the dorsal skull surface of a typical mouse (36) (Figure 1 (b)). This feature aids alignment during attachment, and a large surface area enables a strong adhesive bond to the skull surface. Adhesive cement is applied continuously along all points of contact to create a permanent bond along the entire perimeter of both sides of the chamber (Figure 2 Step 1 (ii)). The cement applied along this joint effectively seals the bottom of the aseptic chamber and is critical for its longterm integrity.

**Fig. 2.**
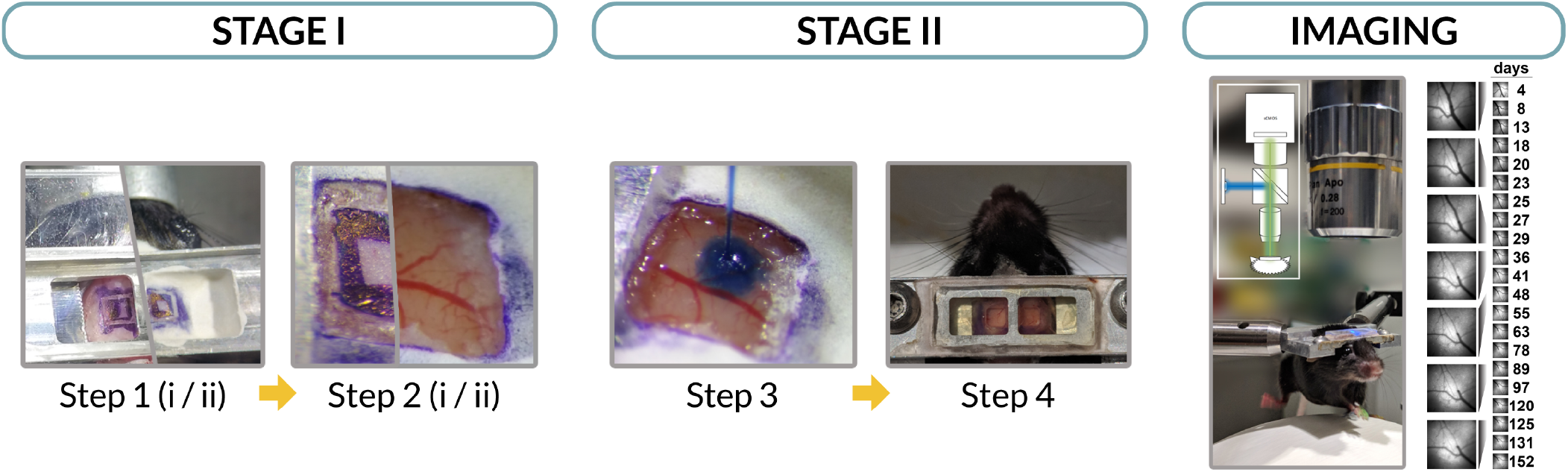
Two-stage surgical installation procedures for the cranial implant. Stage I: headplate attachment to cranium using dental cement followed by bilateral craniotomies. Stage II: this stage can be delayed to allow mouse recovery. If the two stages are performed separately, the exposed tissue can be protected through a temporary silicone seal. It includes experiment-specific tissue interventions, such as injection of virus for cell labeling, and installation of the optical insert. Similarly, the optical insert can be detached from the headplate for future tissue access. Imaging: mouse with attached headplate and cranial window on a spherical treadmill under 10X objective lens of a fluorescence imaging microscope (schematic over photo). At right are sample image frames recorded from the same region over a period of 5 months, beginning 4 days after injection of GCaMP6f virus.

The wide area of skull-to-headplate attachment provides a mechanically stable coupling between the animal’s skull and the headplate holder, which is fixed to the microscope table. The headplate is bonded to all skull plates, which stiffens the skull tremendously. Additional rigidity is provided by a central support structure that contacts the skull along the sagittal suture. All these features combine to provide a very rigid attachment to the mouse cranium, which drastically reduces its motion relative to the imaging system. Remaining brain motion is then primarily movement relative to the skull, and may originate from physiological forces (i.e. cardiorespiratory) as much as behavioral forces from animal movement; suppressing this intracranial motion is addressed in the design of the optical insert described below.

#### Chamber

The chamber in the headplate center facilitates intermittent physical access to neural tissue by protecting the craniotomy sites between tissue interventions (Figure 1 (b)). Once the headplate is bonded to the animal’s skull, the floor of the chamber is formed by the central support structure that traverses and fuses the sagittal suture, the skull surface surrounding each craniotomy, and a flat ledge that extends laterally. The joints between the skull surface edges of the central support, anterior and posterior walls, and the lateral ledge are sealed during the headplate attachment procedure (Figure 2 Step 1 (ii)). This bottom seal is crucial for maintaining an aseptic environment for the protection of the exposed brain tissue. When the dura mater is left intact during the craniotomy, the space within the chamber is continuous with the epidural space.

#### Optical Insert

The insert has optically flat top and bottom rectangular surfaces (Figure 1 (a)). The bottom brainfacing surfaces are positioned to form a flat interface with the intact dura through each craniotomy. The body of the insert provides a clear optical light-path between top and bottom surfaces. The walls of the body are tapered to increase the angle of unimpeded light collection/delivery at the image field. This increases the numerical aperture for imaging through high power lenses, and also expands options for off-axis illumination. The tapered body is extended to the brain surface via vertical-walled columns that traverse each craniotomy (Figure 1 (c)). These columns fill the space made by removal of the bone flap during craniotomy, and their bottom surface gently flattens the brain tissue, positioning the cortex in a horizontal plane for convenient wide-field imaging. Both the top and bottom surfaces are made optically clear by integrating microscope slides in the mold when casting.

Inserts are fabricated in batches using an optically transparent silicone elastomer. We vacuum cast the part in a PTFE and glass mold with an aluminum window frame inclusion that gets embedded near the upper surface (Figure S2). This frame provides a site for attachment and sealing to the rim of the chamber, as well as structural reinforcement. This helps to establish and maintain a flat optical surface at the top of the insert, parallel to the headplate. We constructed inserts with the bottom surface parallel to the top, which works well for imaging medial cortical regions. For imaging lateral cortical regions (e.g. visual or auditory cortex) the mold can be adapted to produce inserts that form a flat image plane with consistent controllable angle relative to the headplate. For any desired angle, this capability greatly simplifies recording from a consistent image plane across sessions and animals. The medial cortical region imaged in the demonstration provided here was square in shape (2 mm X 2 mm), at a horizontal angle of 0 degrees, and extended from 0.83 mm to 2.83 mm symmetrically off the midline.

### Installation and Usage

The surgical installation procedures for this two-stage cranial implant device were adapted from a combination of procedures in common use for attachment of headplates and cranial windows in mice (37, 38), and similar devices used for optical imaging in primates (10, 11, 24). The specific protocol evolved during 3 distinct trial runs, and the final protocol is summarized here and detailed in methods and materials below. The 18 mice reported here received the same version of headplate and optical insert. Minor changes were made to the surgical procedures from one batch to the next, each with discernable effect; see the discussion for details.

Because this is a two-stage cranial implant device, the procedure for installation can be separated into multiple distinct surgeries depending on experimental requirements (Figure 2). The first stage includes headplate attachment to bare skull, centrally aligned along the AP axis with the bilateral sites over the cortical region of interest (Figure 2 (Step 1). Once the headplate is securely bonded, bilateral craniotomy can be made through the skull in the floor of the chamber (Figure 2 (Step 2)). If the second stage of installation is performed separately, the chamber is given a temporary silicone seal to protect the craniotomy. We delayed the second stage of installation for at least 2 to 3 days to allow for mouse recovery.

The second stage involves installation of the optical insert, and may be directly preceded by injection of virus, pharmaceutical compounds, exogenous cells, or any other substance of interest (Figure 2 (Step 3)). The cranial window is installed in the chamber with the assistance of a custom stereotaxic attachment (Figure S3), which enables fine height adjustment and holds the window’s position while being secured in place. The angle of the window’s top surface is held parallel with that of the headplate. The chamber is partially filled with sterile agarose to displace all air from the chamber when the optical insert is lowered into place. The height is adjusted to provide full contact between the insert’s bottom surface and the dura, which also places the insert’s frame in close proximity to the rim of the chamber. Dental cement is applied to form a joint between the headplate and the window frame of the optical insert, installing the insert in place and aseptically sealing the chamber (Figure 2 (Step 4)).

The optical insert can be removed and replaced at any time to provide intermittent physical access to the neural tissue and/or for window replacement (i.e. for mid-experiment injections or window damage repair, respectively). Removal is relatively easy, accomplished by etching away the joint between headplate and optical insert. Window replacement uses the same procedure as the second stage installation described above.

The replacement procedure was attempted 5 times, 4 of which were successful in preserving or restoring optical quality to “like-new” condition, without inflicting detectable tissue damage. Three windows were removed and replaced following damage to the top surface of the optical insert, inflicted by feisty cagemates with sharp incisors (at 91, 83, and 172 days post-installation; 91 days case unsuccessful). The remaining two cranial windows were removed at 20 days post-installation to facilitate direct tissue access for grafting exogenous cells to the imaged region. We found that the removal needs to be performed slowly, taking great care to avoid capillary rupture in the exposed brain and surrounding granulation tissue. During each of these procedures, we observed the pattern of granulation tissue growth into the peripheral area of the chambers (Figure 3 (d)). Photos of the typical growth (as observable with window removed) at day 172 is shown in Figure S5, and described in more detail below.

**Fig. 3.**
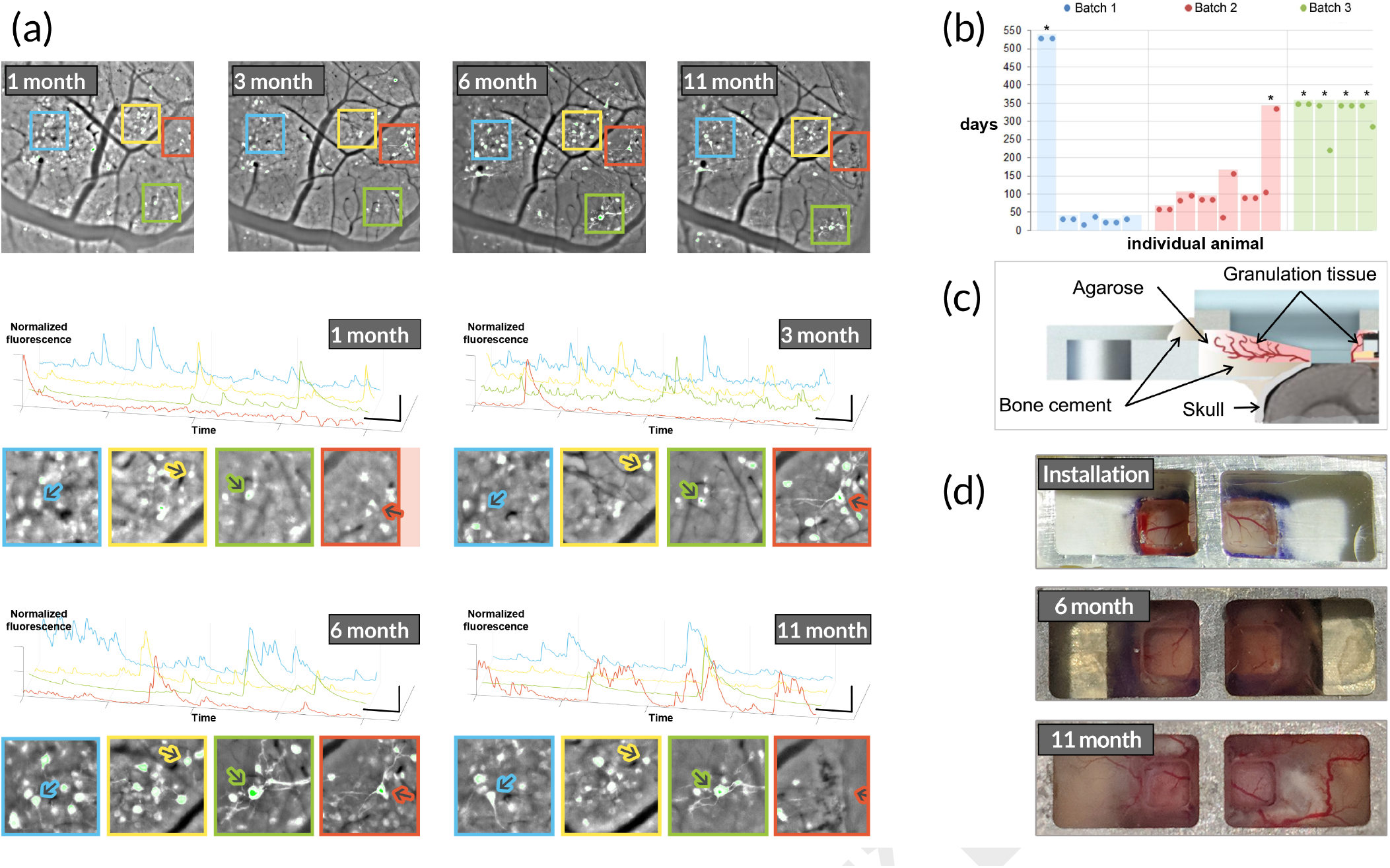
Calcium activity of same cells can be tracked over the period of a year. (a) Long-term optical quality of the window is preserved over a long period to capture calcium activity of neurons over the same imaging region. Selected sessions from a single imaging site show continued tracking of 4 prominent cells over course of experiment. Top: frame extracted from video with prominent cells outlined with colored boxes corresponding to zoom to the bottom. Bottom: Zoom around prominent cells indicated by color and time-series plot of the aggregate pixel intensity of the prominent cells marked with colored arrows in the zoom images to the left. 40 seconds from each session are shown. Pixels are aggregated by adding the intensity from 10 pixels with greatest intensity in each frame drawing from a 32 x 32 pixel rectangle centered on each cell (not shown), then normalized so each time-series stretches between 0 and 1. (b) Summary of batched trials used to evaluate prototype windows as the design evolved. Asterisks are added to indicate windows that persisted greater than 1 year. (c) Cross-sectional schematic of the observed pattern of granulation tissue growth into peripheral areas of the imaging chamber. (d) Granulation tissue growth within the peripheral chamber areas at 0, 6 and 11 months post-installation.

### Evaluation of System Performance

Throughout development we installed several prototypes to test the effect of various features and conditions. The cranial window design and surgical procedures described in this paper were attempted with 18 mice. Cranial window condition was evaluated by direct observation and evaluation of fluorescence dynamics in processed video recorded during periodic 5-minute imaging sessions. Direct (bright-field) observation with a stereoscopic microscope was useful for evaluating quality of the optical interface with brain tissue, as well as for tracking progression of granulation tissue growth in the surrounding space at the edges of the craniotomy. Analysis of cell dynamics measures from processed fluorescence imaging video indicated actual usability of the window for longitudinal studies requiring activity metrics at a cellular level.

#### Experimental Batches

The first batch served as a short trial-run for the prototype and procedures whose performance in early tests suggested strong potential for long-term reliability. We ran the first batch for 4-6 weeks to get a better assessment of what we could expect for long-term viability. With this design and minor modifications to the surgical procedure, we felt comfortable using the window in longitudinal studies with significant investment of time and resources at stake that would also allow for continuous assessment of the window’s optical performance in parallel. The first batch (N = 5) of windows was installed and was evaluated 2-3 times/week for just over 1 month. Several more were installed for use in a long-term imaging study beginning with the second batch (N = 6), then the last batch (N = 5). The results of these runs are reported below, summarized in Figure 3 (b).

#### Sustained Optical Quality Extended over a Year

In the first batch of 7 mice, optical quality provided by the window was more than sufficient to record cell dynamics across both image regions beginning 4 days post-installation and fluorophore-transfection procedure and was sustained for several weeks. At 4–6 weeks post-installation this batch of mice was evaluated and 4 of the 5 mice were discontinued and the installation procedure was adjusted for the next batch. The decision to discontinue in each case was based on observed deterioration in either the health of the mouse (2 out of the 4) or the optical quality of the window (2 out of the 4). See the discussion section for the mechanisms we suspected to underlay and procedural adjustments made to address these issues.

We continued to observe and image the 5th mouse. Progression of the optical quality and fluorophore expression characteristics in bilateral image regions is depicted in Figure 3 (a) for this mouse. Optical quality at the brain-to-window interface has remained consistent for longer than 18 months. The structure of granulation tissue surrounding the window at 11 months is described in detail below and depicted for this mouse in Figure 3 (d).

Similar to the first batch, the second batch of 6 mice was observed and recorded for some time (3-5 months) before discontinuing all except one most exceptional mouse. This mouse received a window replacement at day 83, and was imaged periodically for 11 months before terminating due to a health concern unrelated to the surgical procedure.

The imaging period for the last batch was extended without pre-termination to more thoroughly test the longer-term limits of sustained optical quality. Of 5 mice, 1 mouse did not recover as promptly as expected following the craniotomy procedure and was immediately discontinued. We observed consistent performance on long-term optical quality, extending over 12 months on average among the 8 windows.

#### Direct Observation of the Integrated Chamber

We periodically examined the imaging chamber condition in all implanted mice using a stereoscopic microscope (Figure 3 (d)). Degradation of the optical interface was found frequently in prototypes/procedures that preceded the one mentioned here. This was observed as progressive extension of a cloudy white inhomogeneous layer across the brain-facing surface of the optical insert. Using the design and procedures reported in this paper, this type of degradation rarely occurred, limited to the cases reported above in Batch 1.

Remarkably, but not unexpectedly, tissue growth surrounding the insert was evident in all cases, regardless of window quality. The tissue appeared highly vascularized, and grew from the craniotomy edge outward along the chamber floor (Figure 3 (c)). This growth is a natural response to the tissue damage inflicted by any craniotomy procedure. The difference observed here is only that the growth does not extend under the window. Instead, it forms a non-adhesive interface with the vertical-walled columns and diverges upward into the aseptic chamber, replacing the agarose fill between the optical insert and the adhesive cement covering the skull and chamber floor. To further investigate the structure of granulation tissue growth into the peripheral chamber areas we removed the optical insert for unobstructed observation in several mice. An especially gnarly example from a 6-month duration window is pictured in Figure S5.

### Adaptability

We also report here an adaptation of the more thoroughly-tested headplate and cranial window design described above (Figure 4). The most obvious difference in the newer design is a substantially larger window, which provides optical access to the entire dorsal cortex. Additional features were added to aid window positioning, improve sealing performance, and to simplify fabrication. Adding the additional design features was made possible by switching the headplate fabrication process from CNC machining to 3D printing, as discussed below. Following is a summary of the major changes incorporated in the latest configuration, included here to demonstrate the adaptability of the basic design described above.

**Fig. 4.**
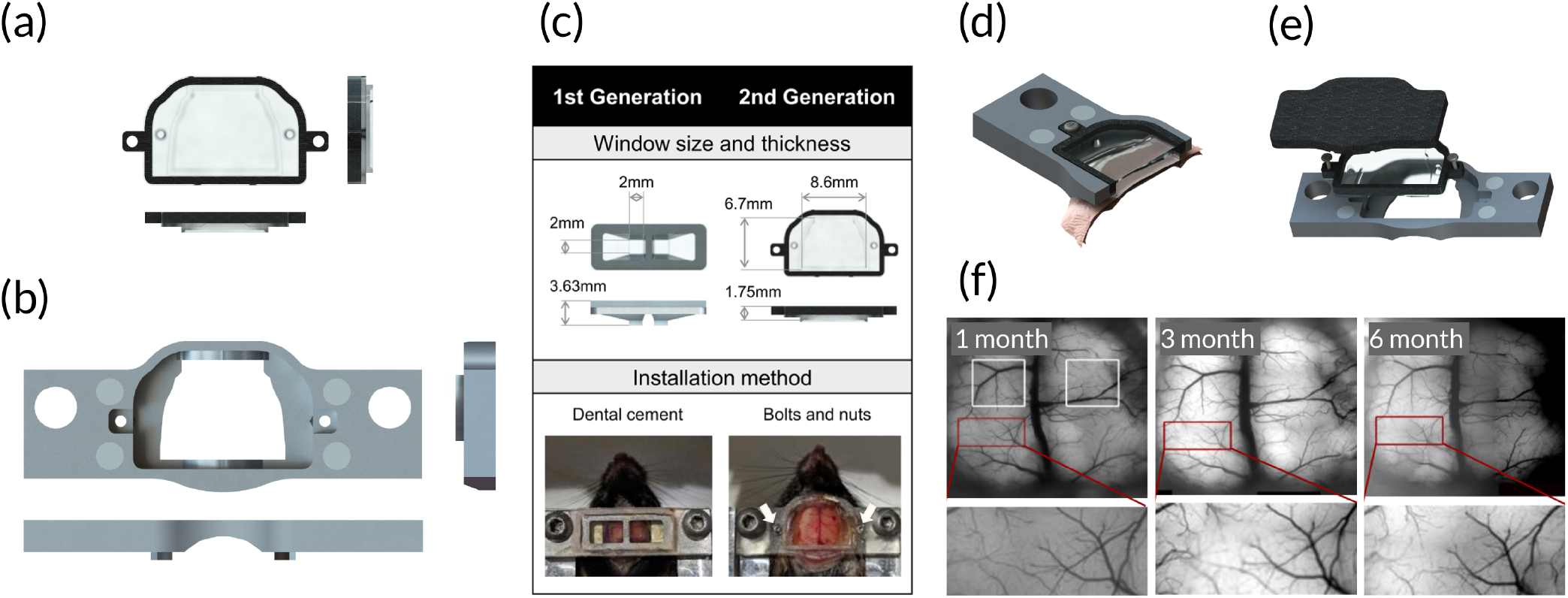
Second generation design of cranial implant system for larger scale cortical imaging. (a) Optical window is expanded for substantially larger imaging area and the thickness is decreased. Tabs with holes are added to the side to provide more precise control over attachment to headplate. (b) Headplate is modified to mate with larger optical window and attach to cranium over a wider base. Protrusions are added to the underside of the plate to stabilize attachment. (c) Comparison of first and second generation designs. (d) Cross-section rendering of model assembly. (e) Exploded rendering of model assembly. (f) Fluorescence imaging frames recorded from a thy1-GCaMP mouse at 1, 3 and 6 months post-installation, demonstrating stability of optical quality over time. The white squares on the 1-month image show the size of the regions imaged with the first generation imaging window for comparison. Red squares indicate the zoomed area below each image.

The height and width of the window frame and chamber were increased, and the window thickness decreased. Protrusions were added to the bottom surface of the headplate which follow the lateral edges of the chamber (Figure 4 (b)). These protrusions contact the mouse cranium along the squamosal suture to maintain a rigid skull-to-headplate attachment despite a reduction in attachment area and removal of a larger fraction of parietal and frontal bone. A thin skirt was added to the bottom surface of the optical insert along its perimeter to help block tissue growth across the imaging area (Figure 4 (a, d)). This is functionally analogous to the vertical-walled column of the prior design, which compensated for the skull thickness, but it accommodates the irregular curves of the endocranium surrounding the larger window. Window installation and height adjustment are improved by fixing small nuts into the headplate on either side of the chamber, and using fine threaded screws to fasten the optical insert in place (Figure 4 (c, d)). This method of window installation provides a vast improvement over the dental cement method used with the prior design and facilitates fine adjustments to the window height. A thin coat of silicone was added to the chamber’s inner walls prior to installation to help seal the upper rim of the chamber with the outer edge of the optical insert. The tape that was used previously to protect the top surface of the implant was replaced with a solid flat magnetically-coupled cap (Figure 4 (e)).

## Discussion

The two-stage cranial implant devices described here were developed to enable reliable long-term optical access and intermittent physical access to mouse neocortex. Our particular application required bilateral cortical windows compatible with wide-field imaging through a fluorescence microscope, and physical access to the underlying tissue for virus-mediated gene delivery and injection of exogenous labeled cells. Optical access was required as soon as possible post-installation, and to be sustained for several months. The design focused on addressing the issue common to other window designs meant for rodents: progressive degradation of the optical light-path at the brain-to-window interface caused by highly scattering tissue growth. The optical insert is molded to fit the chamber and craniotomy, blocking tissue growth and providing a reliable optical interface for up to one year. The core design can be rapidly adapted to improve performance or for a variety of applications.

### Critical Elements

In assessing the design presented here, we can point to a few critical elements that facilitate the maintenance of the long-term optical quality. Refer to the methods section for the specifics of surgical procedures for head-plate attachment and optical insert installation. These procedures were established after testing the variable formulations in protocol (1, 10, 16, 24, 30, 39).

First, the design of the optical insert must incorporate a mechanical barrier that fits along the edges of the craniotomy. The barrier must be continuous along the circumference, and extend as far as the inside surface of the skull to be effective. Achieving this tight fit without aggressively impinging on the brain requires some sort of fine height adjustment capability. The optical insert must be installed at the correct height during the installation procedure, or shortly thereafter. The insert must be depressed very slightly until full contact is made across the entire window, but pressing beyond necessary will quickly exert an undesired increase in intracranial pressure, increasing inflammation and adverse outcomes. The mechanism for fine adjustment can be designed into the system, as is demonstrated in the second design presented here, or incorporated into the installation procedure, as is done in the first design.

Of particular note, we found that administration of antibiotic and anti-inflammatory drugs in the days surrounding any major surgical procedure had a substantial impact on the viability of the optical interface. We used both corticosteroid and non-steroidal anti-inflammatory drugs, and attempts to exclude either ended poorly more often than not.

Equally critical to the long-term health of the imaging chamber was the requirement to establish and maintain an air-tight seal between the chamber and the outside world. This includes a permanent bond between the chamber and skull, and a reversible bond between the chamber rim and optical insert. How this is accomplished will be specific to the system design, but it is absolutely essential.

In addition to establishing and maintaining an air-tight seal, it is necessary to eliminate any and all pockets of air within the chamber. Any air pockets that remain after installation will be susceptible to bacteria growth and may disrupt normal intracranial and intermembrane pressures. The system presented here used sterile agarose fill to displace all air within the chamber prior to sealing. Dead space surrounding the optical insert, including that temporarily filled with agarose, will fill with fluid and eventually be overtaken by granulation tissue. This process is helpful to the maintenance of a aseptic chamber environment, so care should be taken not to disrupt it. However, an excess of dead space will delay this process, and thus should be minimized when adapting the design.

Many attempts to test variations from the described procedures indicated that all elements mentioned above are equally critical to achieving a reliable imaging window with sustained optical quality. Implementing the procedures as described or something similar should mitigate the primary obstacle to long-term imaging in mice and other rodents. The need to pre-terminate imaging experiments due to optical light-path disruption by tissue ingrowth should be substantially reduced.

### Staging Implant Installation and Tissue Access

Configuring the implant as described, so as to enable a staged installation of multiple parts enables surgical procedures to be spread across multiple days. This capability offers a number of advantages. It may save time and resources - particularly during the prototype stages - by allowing time to ensure each implanted mouse fully recovers from the initial procedure. Additionally, the delay between surgeries allows the heightened inflammation and other immune system response triggered by craniotomy to normalize before attempting a tissue intervention that is sensitive to these conditions (e.g. viral or cell injections). Through this mechanism the system offers the capability to image the first tissue intervention from day 0.

Similarly, designing the system to be installable in multiple stages enables trivial and repeatable tissue access at later time points by simple reversal of procedure for optical insert installation. The process may be comparable to a previously reported method of removing the entire glass window to access the tissue (16). With this system, however, the methods used to remove and replace are faster and simpler and carry less risk of tissue damage compared. Additionally, the described methods of facilitating tissue access can be advantageous over a fixed access port by providing full access without compromising the image field (12–14).

### Design Adaptation

While the specific designs described in this report have much to offer, the greatest asset of the underlying system is its easy adaptability. The design can be rapidly transformed to accommodate various applications or to modify its performance in response to new technologies and demands. This rapid adaptability was a primary goal of this project, and informed our design and engineering decisions throughout development. Anyone with access to common laboratory equipment and moderate engineering and fabrication skills can produce a system to fit their particular needs. As an inherent aspect of any design process, the adaptation of the original design evolved over the course of prototyping and testing (Figure S7). In presenting two designs in this report, our intention was to demonstrate the technical feasibility of continuous development of a futureproof system. The original system was adapted to accommodate the continuous evolution of image sensor technology, particularly the growth in size and resolution, expanding the field of view and allowing simultaneous access to cellular interactions across multiple brain regions using wide-field imaging (7, 33, 34).

The iterative process used here was only made possible by using the now widely available rapid prototyping procedures, 3D-printing and lasercutting – major progress of manufacturing and its increased versatility, providing better quality, customization, lower cost and shorter production time (40). In an effort to compare various manufacturing technologies, we explored manufacturing the finalized product design through a number of companies and advanced with 3D metal printing with overall satisfaction at i.materialise (Leuven, Belgium) – we had also developed the parts through other rapid prototyping companies including Shapeways (New York, NY) and Sculpteo (Villejuif, France).

## Methods and Materials

### Device development and fabrication

Components were designed using SolidWorks. Prototypes were fabricated using CamBam to generate toolpaths in G-code for machining on a CNC mill. The headplate and window frame were milled from aluminum plate. The mold for casting optical inserts was designed in three parts (Figure S2). The middle component was milled from PTFE. The outer components were made using a laser-cutter and acrylic sheet. CAD files are shared on (https://github.com/susiescha/cranial-window-models.git).

### Window casting procedure

The optical inserts were fabricated through a vacuum casting procedure (Figure S2). Prior to casting, window frames and two glass coverslips (Corning, 2947-75×38, Corning, NY), prepared in advance through plasma etching for 30 seconds and silanization using Trichlorosilane (Sigma-Aldrich, 448931-10G, St. Louis, MO), were inserted into the mold. The mold was then placed between two custom-made acrylic plates with silicone gaskets in between and was assembled using bolts around the perimeter. The pressure control port (McMaster-Carr, 5454K61, Elmhurst, IL) was connected to the house vacuum line, and the fill port (McMaster-Carr, 2844K11) was connected to uncured PDMS polymer (Dow Corning Sylgard (1:10 by weight), thoroughly mixed and degassed in advance. The liquid-state polymer was then drawn into the mold filling the volume in between the two coverslips using vacuum. Once polymer displaced all air, vacuum was released and positive pressure was applied through the pressure control port after plugging the fill port. While maintaining positive pressure, the polymer was cured at 75ºC for 12 hours. Finally, the windows were released from the mold and trimmed using scalpels. Windows were handled so as to protect the top and bottom surfaces from damage or debris. The completed window was sterilized in an autoclave before use.

### Surgical procedures

Animal care for surgical procedures is described below, and the details specific to each procedure are in the sections that follow. All procedures were approved by the Institutional Animal Care and Use at Boston University. Stereotaxic surgeries were performed on 6 to 8 weeks old female C57BL/6 mice (Charles River Laboratories, Wilmington, MA). Pre-operative care for the initial headplate and craniotomy procedure included subcutaneous administration of meloxicam (NSAID, 2.5 ug/g) and buprenorphine (opioid analgesics, 0.3 ug/g), and intramuscular injection of dexamethasone (corticosteroid, 5 ug/g) one hour before surgery. Meloxicam and buprenorphine were continued postoperatively every 12 hours for 48 hours. Meloxicam was also given before and after procedures where brain tissue was exposed, i.e. those for viral or cell injections and window replacement. For all procedures described below, mice were placed under general anesthesia with isoflurane mixed with oxygen.

### Headplate attachment and craniotomy

We shaved the top of the mouse’s head and sterilized the skin using 70% alcohol and 7.5% Betadine. We made a 1 cm midline sagittal incision through the scalp using surgical scissors, and retracted laterally using a self-retaining retractor (WPI, 501968, Sarasota, FL). To prepare the cranial surface, we applied 3% hydrogen peroxide to oxidize and facilitate removal of periosteal tissue with cotton tip swabs. The surface was then marked up before headplate attachment followed by craniotomy. We targeted laterally symmetric craniotomies with edge length 2.2 mm centered at coordinates ± 1.83 mm lateral to sagittal suture and 1.00 mm anterior to bregma. First, we used a surgical skin marker (FST, 18000-30, Foster City, CA) to roughly indicate the site of each craniotomy and enhance contrast of the edges to be etched (Figure 2 (Step 1 (i))). We etched the corners and edges using a dental drill with a FG 1/4 round carbide burr. This way of marking the edges facilitates headplate placement and also aids recovery of the intended craniotomy position despite being covered by semi-transparent adhesive cement in the following steps.

We used a custom stereotaxic attachment to position the headplate symmetrically aligned with the marked sites (Figure S3), and to hold it horizontal while bonding to skull. The headplate was anchored directly to the skull using either opaque or semi-clear quick adhesive cement (Parkell, S380, Brentwood, NY) (Figure 2 (Step 1 (ii))). Subsequently, we began each craniotomy by drilling along the marked edges (Figure 2 (Step 2 (i))). We frequently stopped to flush debris from the site using sterile saline and an aspirator. Once separated from the surrounding skull, the bone flap was carefully removed using a pair of sharp forceps (FST, 91150-20) and a 45º micro probe (FST, 10066-15) while keeping the dura intact (Figure 2 (Step 2 (ii))). At this point, we either attached the optical insert or sealed the area with a layer of non-toxic silicone adhesive (WPI, KWIK-SIL).

### Optical insert installation

The optical insert installation can be performed immediately following the craniotomy or deferred to the day of injection as described below. First, we filled the chamber with sterile 0.5% agarose solution, immersing the exposed brain. Enough agarose was added so as to overflow the walls of the chamber as the window is inserted, ensuring no air gaps remain in the space between the walls of the chamber and the window, below the joint to be sealed. Next, the window was placed in the chamber, directly over the craniotomy, in gentle contact with the exposed tissue. We used a custom stereotaxic attachment to adjust the window height and secure its position during installation. This was aided by an attachment – similar to that used for headplate attachment – which fixed the angle of the window’s top surface parallel with that of the headplate. The height adjustment required depressing the window until full contact was observed between the inner window surface and the dura. The window was tacked in place by applying an accelerated light-cured composite (Pentron Clinic, Flow-It ALC, Wallingford, CT) in at least three points, bonding the window frame to the anterior and posterior walls of the head-plate. At this point the guide was removed and the joint was prepped for sealing. Excess agarose (polymerized overflow from the window insertion step) was aspirated away to expose and clean the headplate surface surrounding the window. The chamber was sealed by filling the joint between headplate and optical insert with dental cement (Stoelting, 51458, Wood Dale, IL) using a P200 pipette. The window surface was protected by applying a double layer adhesive strip made of gaffers tape over a transparent adhesive film dressing (3M: Tegaderm, 70200749201, Maplewood, MN).

### Window removal and replacement

The dental cement connecting the window and headplate was etched away using a dental drill. Before removing the window, we thoroughly flushed debris from the surrounding surfaces with sterile saline. Replacement windows were installed using the same procedures described above for initial installation. Localizing the replacement window to the same position was aided by the expansion of granulation tissue up to the vertical-walled columns of the prior window.

### Injection

The exposed brain was flushed with sterile saline before and after each injection. Injections were made using pulled glass micropipettes with inner tip diameter ranging from 40 and 80 um (WPI, 504949). The micropipette was initially back-filled with mineral oil, then secured onto a microprocessor controlled injector (WPI, NANOLITER2010). The micropipette was then frontloaded with virus or cells using a controller (WPI, Sys-Micro4). In general, an injection of 230 nL of cells labeled with AAV9.CAG.GCaMP6f (AV-9-PV3081, Penn Vector Core, Philadelphia, PA) at 106 cells/uL, or 230 nL of AAV9.Syn.GCaMP6f (AV-9-PV2822, Penn Vector Core). Injection was performed approximately 500 um deep into the cortex at the rate of 46 nL/min near the center of the imaging field, while avoiding blood vessels to maximize the observable cells around the injection site. The micropipette was left to sit for an additional 2 min at the injection site before slow withdrawal.

### Wide-field in vivo imaging and microscope setup

Wide-field epifluorescence imaging was accomplished using a custom microscope equipped with a sCMOS camera (Hamamatsu, ORCA Flash 4.0 V3, Shizuoka, Japan), 470 nm LED (Thorlabs, M470L3, Newton, NJ), excitation and emission filters of 470/25nm banpass and 525/50nm bandpass, a dichroic mirror (495nm), and a 10x objective lens (Mitutoyo, 378-803-3, Kanagawa, Japan). Mice were positioned under the microscope for imaging using a custom headplate holder (Figure S4) and allowed to run on an air-supported spherical treadmill (26). The camera recorded a field-of-view of approximately 1.3 mm × 1.3 mm using an image resolution of 2048 × 2048 or more commonly 1024 × 1024. Continuous image sequences were acquired at 40 to 60 frames-per-second for 5 to 7 minutes. We selected the field to image within each site by roughly centering around the injection site. To focus the microscope on labeled cells in the superficial layers of cortex, we focused on the surface vasculature to find a stable reference, then advanced the focal plane 50 to 150 um until multiple cells were distinguishable. A reference image of the selected image was recorded for each site and used later to reacquire the same field across image sessions. Alignment to this reference image relied primarily on using the major blood vessels as landmarks to guide microscope position in the XY plane. Image sequences were stored for subsequent processing and analysis.

---

**Supplementary Note 1: Model drawings with dimensions in wireframe**

**Fig. S1.**
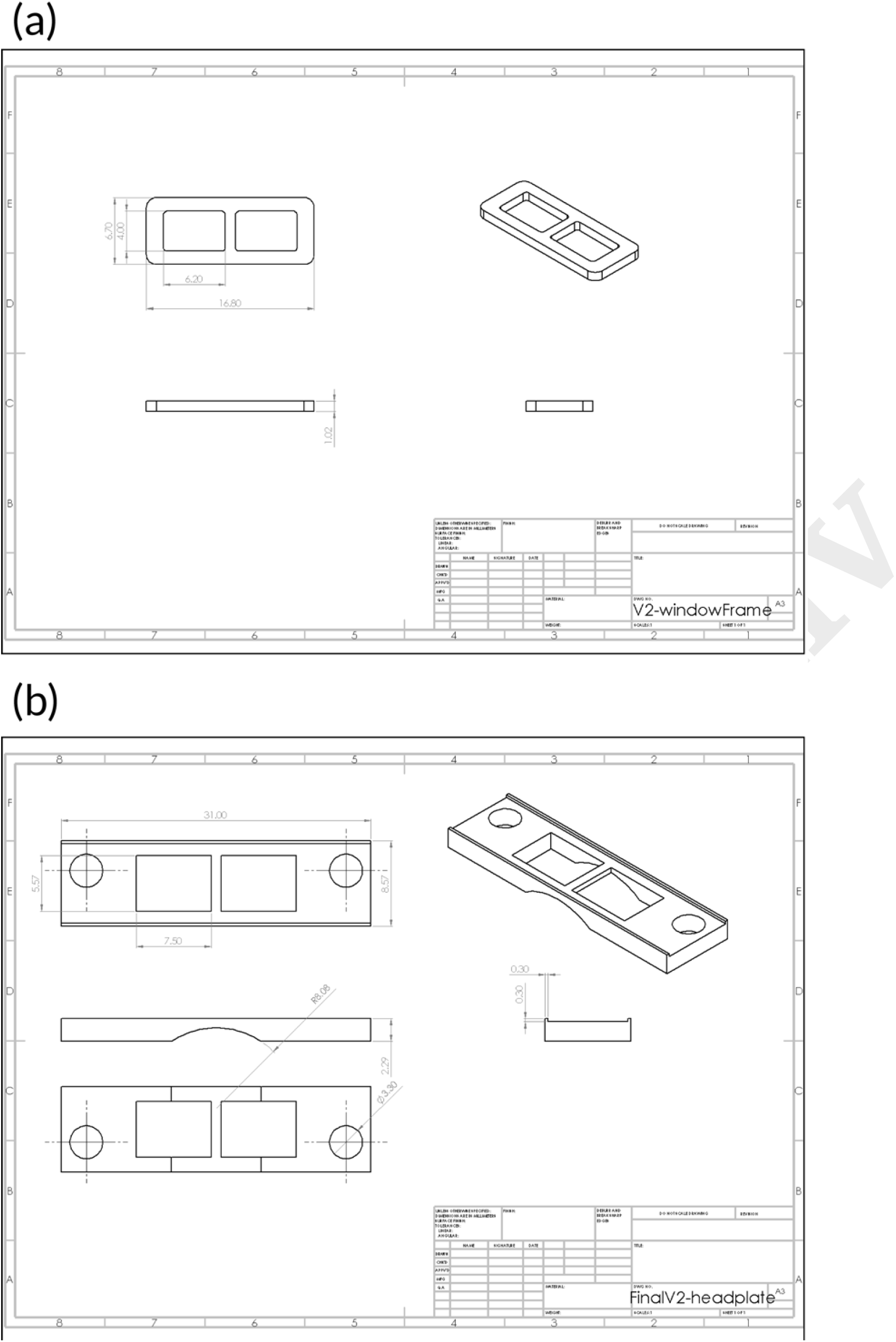
Multi-component cranial window system: model drawings with dimensions in wireframe. Window frame component of optical insert (a). This piece is milled or stamped out of aluminum sheet and gets embedded near the upper surface of the optical insert during casting. This frame provides a site for attachment and sealing to the rim of the chamber, as well as structural reinforcement. Headplate (b).

**Supplementary Note 2: Vacuum casting assembly**

**Fig. S2.**
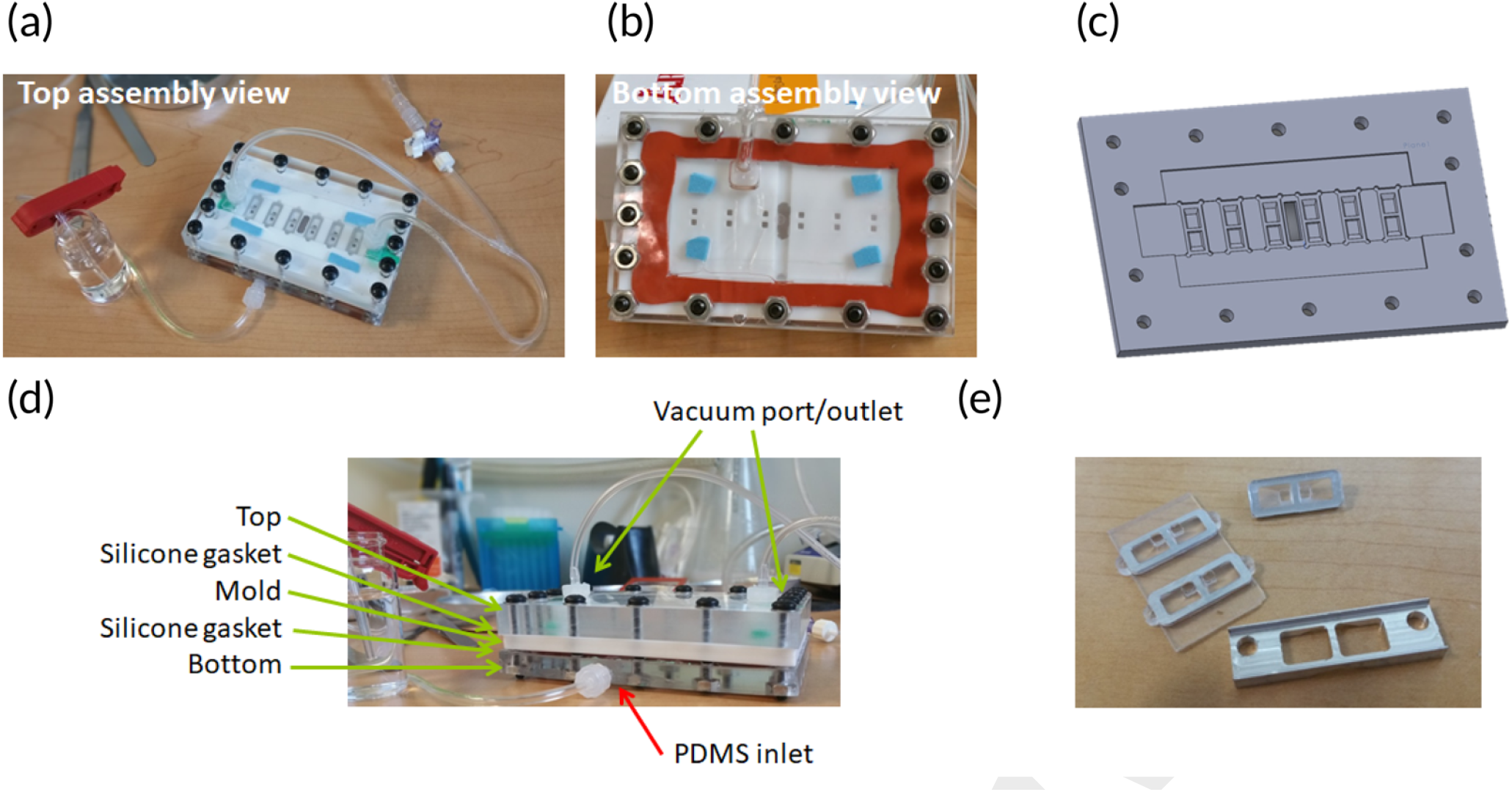
Vacuum casting assembly: (a) Top of assembly, shown during casting. (b) Bottom of assembly, showing PDMS fill-port/inlet. (c) Rendered model of middle mold component. This piece is fabricated on a CNC mill out of PTFE. (d) Side view of assembly; PTFE mold placed between two laser-cut acrylic plates. (e) Optical window pieces after casting with PDMS, before and after separating and trimming the individual windows.

**Supplementary Note 3: Model drawings of surgery guide used during headplate installation**

**Fig. S3.**
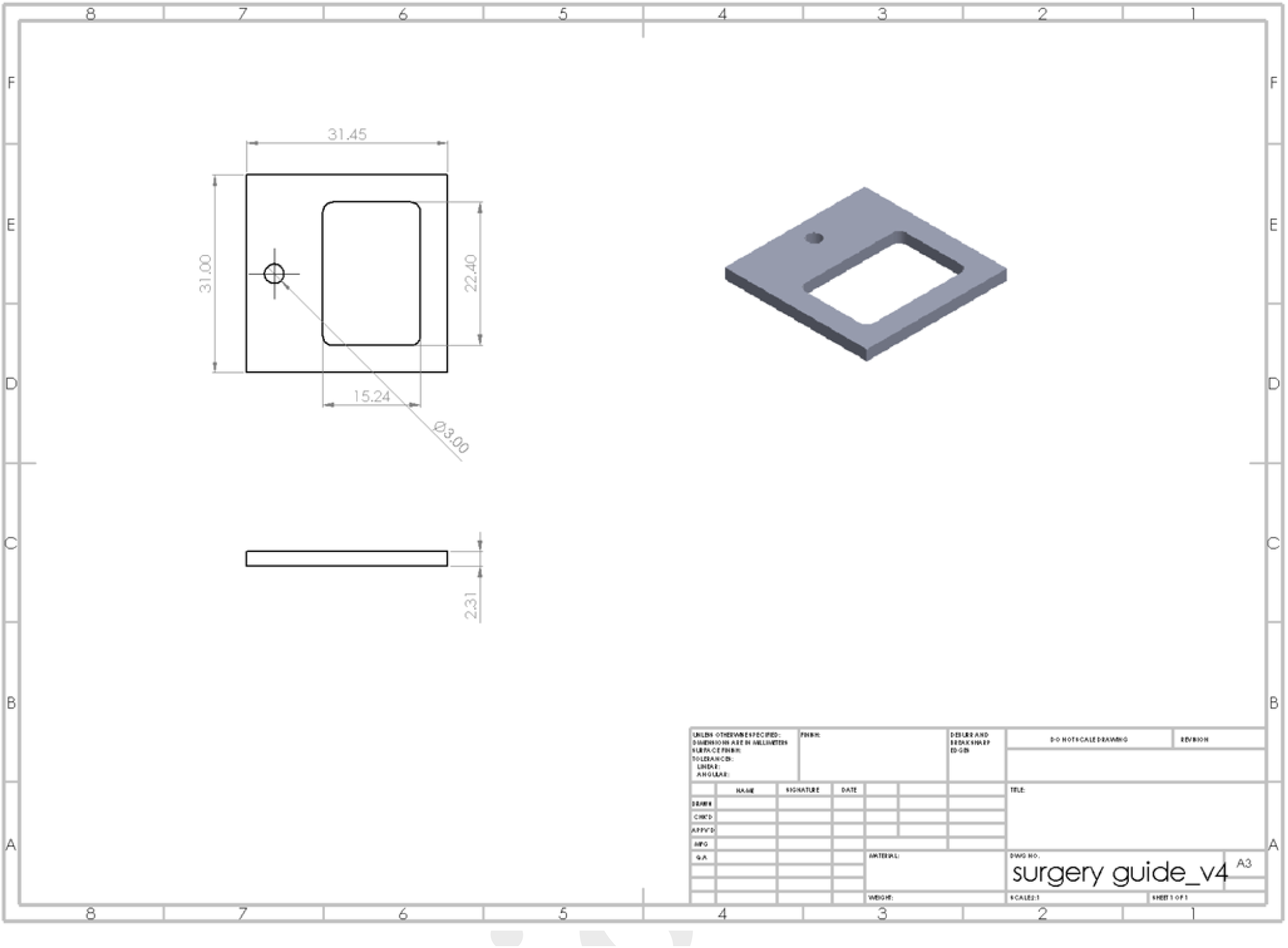
Model drawings of surgery guide used during headplate installation. This component is attached to a common stereotaxic device and fixes the angle of the headplate during attachment, for consistency with a horizontal imaging plane on the microscope.

**Supplementary Note 4: Rendered model of headplate holder**

**Fig. S4.**
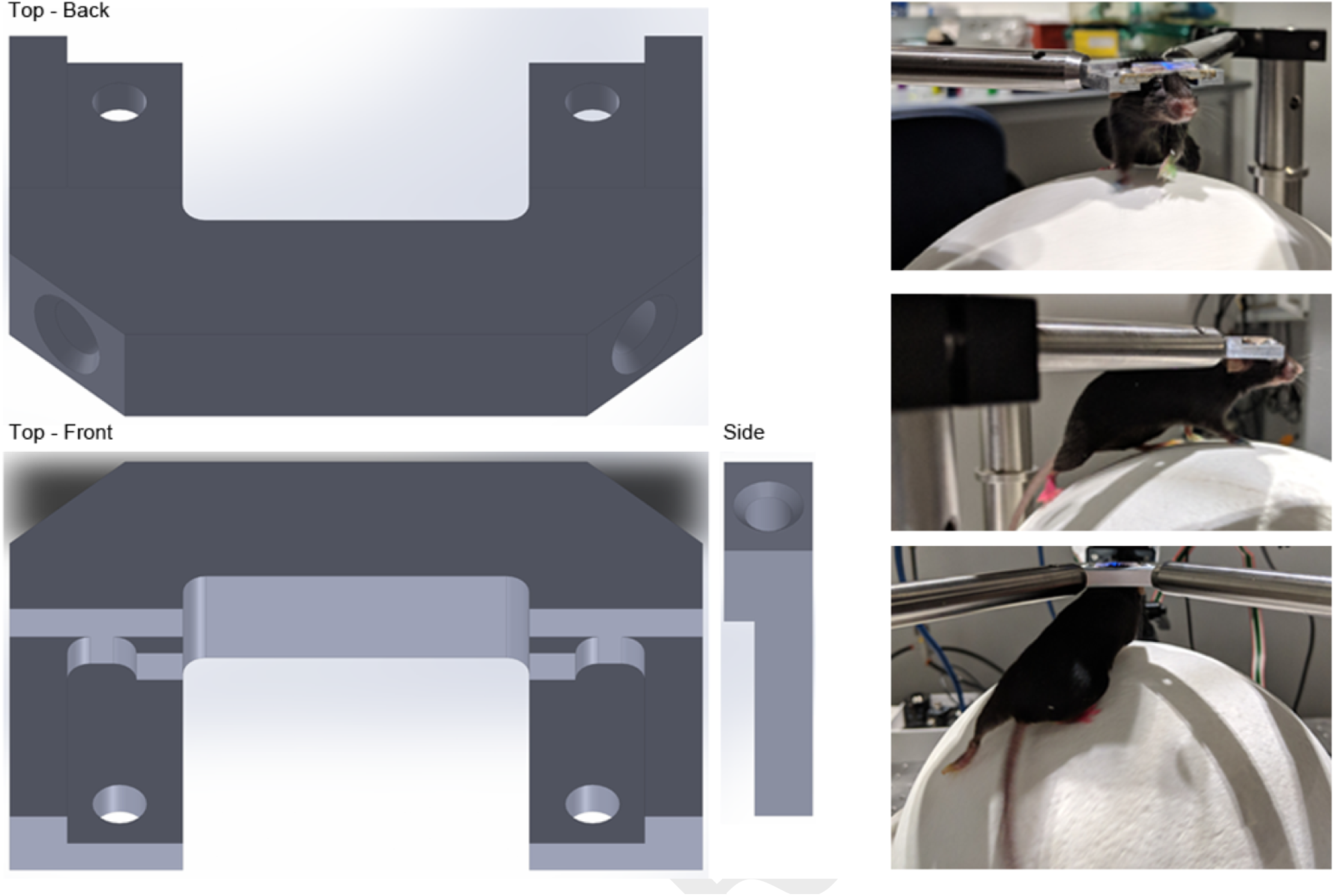
Rendered model of headplate holder: several features of the design are included to simplify fabrication on a benchtop CNC mill by minimizing re-fixturing and tool-changes. The headplate attaches using two M3 screws spaced 1-inch apart (0.5 inches off center). Threaded holes on the angled rear faces of the holder for easy attachment to standard 1/2-inch diameter optical posts. The critical functionality of this component is providing rigid attachment of the headplate to the optical table on which the microscope is built, minimizing relative motion. This part can be fabricated out of aluminum if plastic headplates are used, but if the headplate is made of aluminum it is best to make it out of another metal such as brass, steel, or titanium.

**Supplementary Note 5: Photos of granulation tissue growth**

**Fig. S5.**
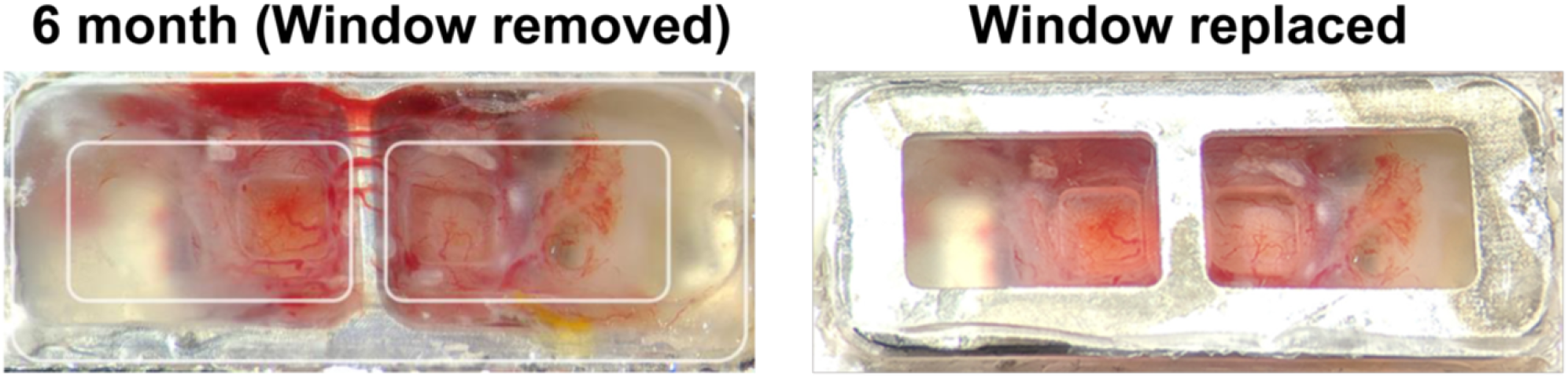
Photos showing growth of highly vascularized granulation tissue into the imaging chamber periphery at 6 months post-installation with window removed (left) and replaced with a new window (right).

**Supplementary Note 6: Model drawings of headplate, window-frame, and magnetic cap of the second-generation system**

**Fig. S6.**
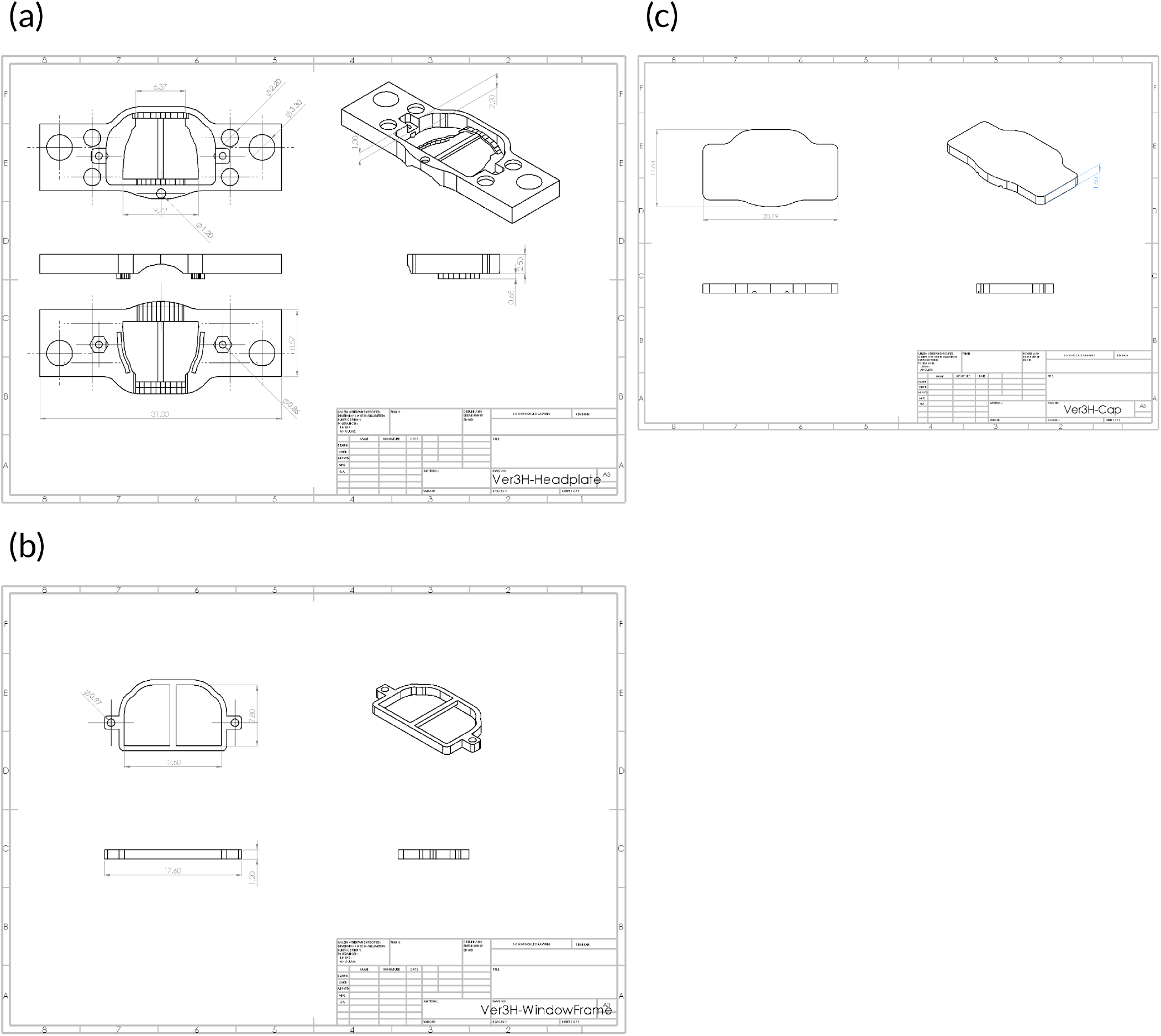
Model drawings of headplate (a), window-frame (b), and magnetic cap (c) of the second-generation window system. The midline bar in the window can be retained or easily removed depending on experiment requirements.

**Supplementary Note 7: Generation of designs**

**Fig. S7.**
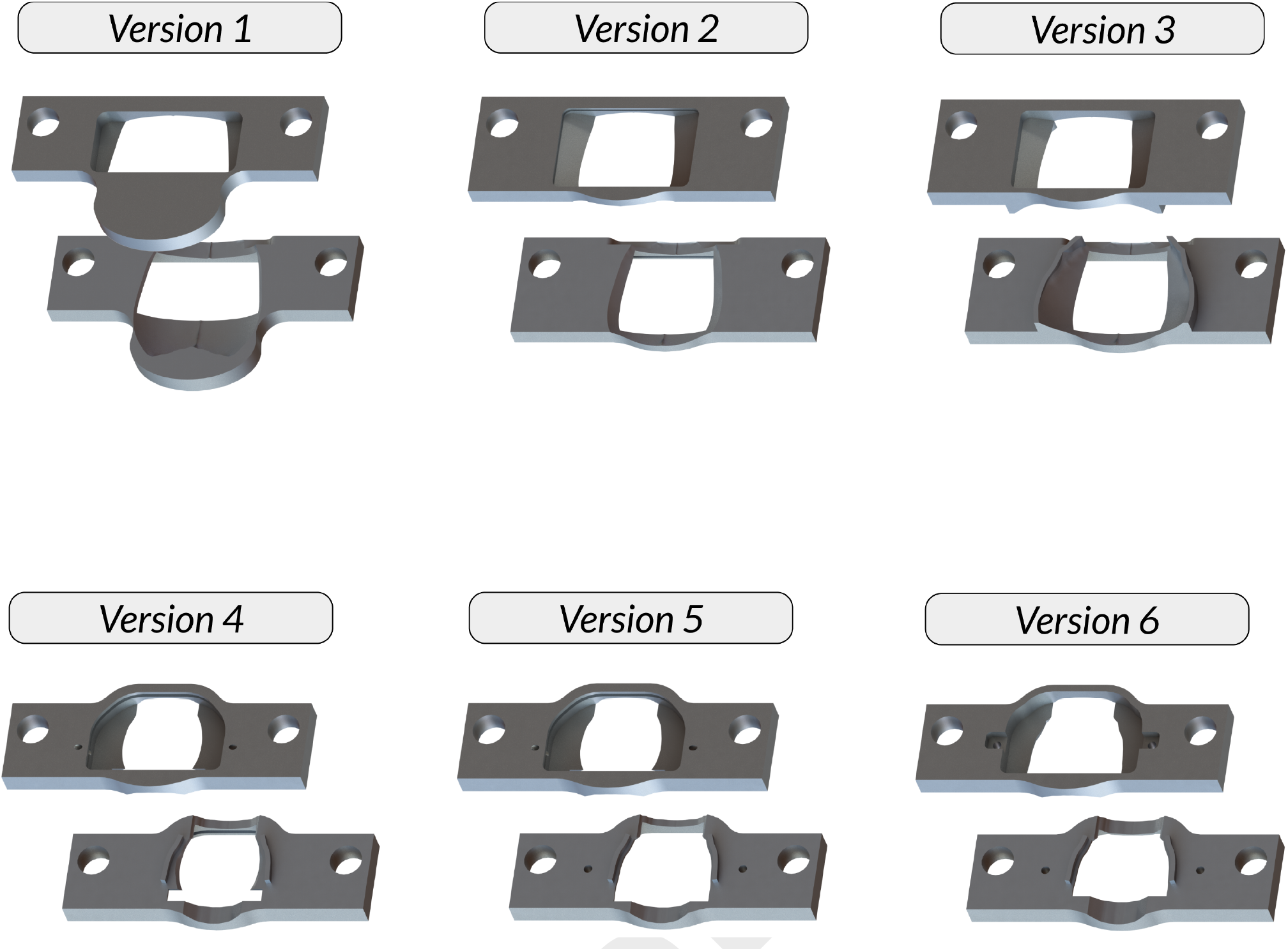
Generation of designs. Additive manufacturing (3D printing – metal, hybrid), laser cutting and CNC can greatly facilitate the prototyping processes.

